# Prior Fc Receptor activation primes macrophages for increased sensitivity to IgG via long term and short term mechanisms

**DOI:** 10.1101/2023.11.14.567059

**Authors:** Annalise Bond, Sareen Fiaz, Kirstin R Rollins, Jazz Elaiza Q Nario, Samuel J Rosen, Alyssa Granados, Maxwell Z Wilson, Meghan A Morrissey

## Abstract

Macrophages measure the ‘eat-me’ signal IgG to identify targets for phagocytosis. We wondered if prior encounters with IgG influence macrophage appetite. IgG is recognized by the Fc Receptor. To temporally control Fc Receptor activation, we engineered an Fc Receptor that is activated by light-induced oligomerization of Cry2, triggering phagocytosis. Using this tool, we demonstrate that Fc Receptor activation primes macrophages to be more sensitive to IgG in future encounters. Macrophages that have previously experienced Fc Receptor activation eat more IgG-bound cancer cells. Increased phagocytosis occurs by two discrete mechanisms – a short- and long-term priming. Long term priming requires new protein synthesis and Erk activity. Short term priming does not require new protein synthesis and correlates with an increase in Fc Receptor mobility. Our work demonstrates that IgG primes macrophages for increased phagocytosis, suggesting that therapeutic antibodies may become more effective after initial priming doses.

## Introduction

Macrophages eat pathogens and infected, cancerous or dying cells via phagocytosis. To select targets for phagocytosis, macrophages measure ‘eat-me’ signals, like IgG antibodies. IgG is recognized by the Fc Receptor (FcR), which is phosphorylated and recruits the kinase Syk, triggering downstream signaling^1,2^. Therapeutic IgGs like Rituximab or Trastuzumab trigger Antibody-dependent Cellular Phagocytosis (ADCP) or Antibody-dependent Cellular Cytotoxicity (ADCC) to reduce cancer growth^2–7^. Even many antibodies originally designed to block the function of their target actually activate the FcR for full efficacy^8,9^. Given the therapeutic importance, there is substantial interest in understanding how to boost macrophage phagocytosis.

What affects macrophage appetite? One important parameter is how sensitive a macrophage is to ‘eat-me’ signals. Antibody-dependent phagocytosis requires the coordinated activation of a sufficient number of FcRs^10^. Targets with a subthreshold amount of IgG are not phagocytosed, despite triggering the initial steps in the phagocytosis signaling pathway^10^. In other macrophage signaling pathways, low levels of activating signal do not elicit any response on their own, but prime macrophages for rapid and intense response to future stimuli^11^. Whether subthreshold FcR signaling has any effect on macrophage appetite is not clear.

During an immune response or treatment with a therapeutic antibody, macrophages encounter multiple potential targets for phagocytosis sequentially, leading to bursts of FcR activation. Some encounters with antibody-opsonized cells result in phagocytosis of the entire cell^3,5^, but many cells do not have sufficient antibodies to trigger phagocytosis. Instead macrophages may trogocytose, or nibble, a target cell or simply ignore it^12–15^. In some circumstances, prior phagocytosis increases macrophage appetite^16,17^. In contrast, other studies demonstrated that phagocytosing several whole cancer cells reduces macrophage appetite^18^. There is no clear, unifying model explaining these differences, which could be dependent on the specific ‘eat-me’ signal presented, the time since phagocytosis, the intensity of the ‘eat-me’ signal, digestion of the internalized particle, or any number of other factors^19^.

To unravel how prior IgG exposure affects macrophage appetite, we need to precisely control the timing and intensity of activating specific phagocytic receptors. Delivering a temporally controlled, homogenous antibody stimuli to a population of cells is very difficult with the current tools. Because soluble IgG does not activate the FcR, IgG must be presented on antibody-bound targets. Controlling precisely the number of targets a macrophage encounters, or the timing of these encounters is very difficult and low-throughput.

To quantitatively control the duration and strength of FcR activation, we developed an optogenetic FcR (optoFcR). We found that prior FcR activation primes macrophages for greater responses to subsequent stimuli. Counterintuitively, low levels of optoFcR activation induced stronger priming than high levels of optoFcR activation. Macrophage priming is controlled by two independent mechanisms, one short-term (<1 hour) and one long-term response (starting at 4 hours, and lasting up to 3 days). The short term response is associated with an increase in FcR mobility, accelerated initiation of phagocytosis and increased phagocytic cup formation. The long term response requires activation of Erk to drive a transcriptional response. These data suggest that macrophages can integrate signaling from previous encounters with IgG to modify the response to the next target. This study provides insight into how macrophage appetite is regulated, and may suggest strategies to enhance antibody-dependent cellular phagocytosis.

## Results

### Optogenetic FcR recapitulates native FcR signaling for precise temporal control over signaling

To precisely control the temporal pattern of FcR activation across an entire field of cells, we sought to design an optogenetic FcR that could be turned on and off with light. Prior work has shown that the FcR clusters upon IgG binding. Although the FcR has no inherent kinase activity, clustering promotes phosphorylation of the intracellular Immunoreceptor Tyrosine-based Activation Motifs (ITAMs) and phagocytosis^20–22^. We hypothesized that clustering may be sufficient to induce FcR activation. To test this, we designed an optoFcR construct that consists of a myristoylation sequence for membrane localization, the intracellular domain for the native FcR gamma-chain and a light activatable peptide CRY2 (Figure 1a). We selected the variant CRY2olig because it forms multimeric, rapidly reversible clusters^23^. Upon blue light (450nm) stimulation the optoFcR oligomerizes (Figure 1b; Video S1). Clusters dissociate when the cells are returned to the dark (Figure 1b). Clustering also results in the recruitment of the downstream effector protein Syk, suggesting that clustering is sufficient to induce FcR phosphorylation (Figure 1c,d; Video S2).

**Figure 1.**
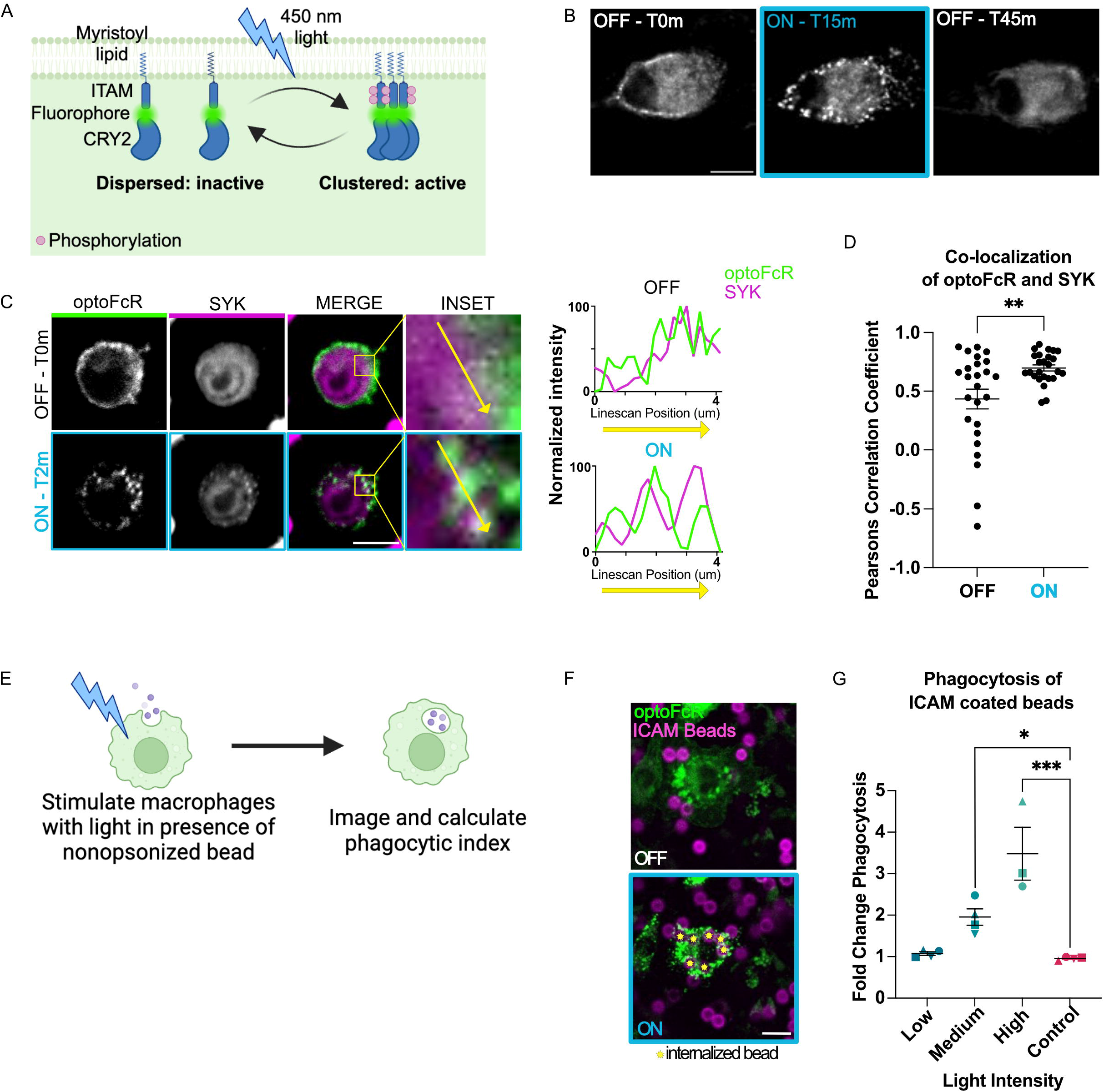
Optogenetic Fc Receptor (optoFcR) controls phagocytosis. A) Schematic of optoFcR design, containing a myristoylation signal for membrane localization, the ITAM-containing intracellular domain from the FcR gamma-chain for signaling, iRFP fluorophore, and the homo-oligomerizing peptide - cryptochrome 2 (CRY2olig). B) Representative images of optoFcR in bone marrow derived macrophages (BMDMs). The optoFcR starts dispersed on the membrane (T0m; n=0/22 cells with visible clusters), and clusters after 15 minutes of constant high intensity light (T15m; n=21/22 cells with visible clusters). The optoFcR is declustered after 30 minutes in the dark (T45m; n=0/22 cells with visible clusters). See also supplemental movie 1. C) Representative images of downstream effector protein, SYK, colocalized with optoFcR in Raw macrophages stably expressing both the optoFcR and SYK-mNeonGreen. On the right, a line scan shows the Syk and optoFcR intensity at the position indicated by the yellow arrow in the inset. See also supplemental movie 2. D) Graph shows the Pearson’s correlation coefficient for Syk and optoFcR at the cell cortex. Each data point represents a cell with an ROI drawn around a membrane region. The same ROI was used for both T0 (OFF) and T2 (ON). E) Schematic of experimental design for f-g. F) Representative images show optoFcR (green) expressing macrophages and ICAM conjugated beads visualized by atto390 in the supported lipid bilayer (magenta) before (OFF) and after (ON) 15 min of optoFcR activation with high intensity light. Internalized beads are labeled with a yellow star. See also supplemental movie 3. G) Quantification of phagocytosis in BMDMs after 15 minutes of optoFcR stimulation at low (5 uW/cm^2^), medium (90 uW/cm^2^), and high (1390 uW/cm^2^) intensity light compared to control cells that do not express the optoFcR but receive the high intensity light stimulus. Both medium and high intensity light stimulate phagocytosis. Each data point represents the mean of an independent experiment. Data collected in the same replicate are denoted by symbol shape. In all graphs, bars represent the mean and SEM. * indicates p<0.05, *** indicates p<0.0005 using a paired t-test (d), one way ANOVA with dunnett correction (g). Scale bars are 10 um. See also supplemental figure 1.

We next sought to determine if clustering of the optoFcR is sufficient to trigger phagocytosis. We incubated ICAM-1 conjugated beads with macrophages expressing either the optoFcR or membrane tethered mCh (mCh-CAAX) and exposed them to 15 min of light stimulation (Figure 1e). ICAM-1 mediates bead binding to the macrophage, but does not trigger phagocytosis of otherwise unopsonized beads^24,25^. Macrophages expressing the optoFcR engulfed three times as many beads as control macrophages when stimulated with the highest intensity light and twice as many beads when stimulated with medium intensity light (Figure 1f,g; Figure S1; Video S3). Low intensity light did not activate phagocytosis. This dose response is similar to the dose response seen in IgG mediated phagocytosis (Figure S1). Together, these data demonstrate that clustering of the FcR ITAM domain is sufficient to initiate phagocytosis in macrophages without a specific ‘eat-me’ signal.

### Prior FcR activation enhances phagocytosis of IgG coated beads

With the ability to temporally control FcR activation in bone marrow-derived macrophages, we next sought to determine if prior FcR activation influences phagocytic ability. We stimulated macrophages with low intensity blue light to activate the optoFcR for 15 minutes which is around the same timescale that a macrophage interacts with a phagocytic target. We then waited 1 or 12 hours before adding IgG opsonized beads, which is sufficient time for the optoFcR to completely decluster. Then we measured the number of beads engulfed per cell using microscopy (Figure 2a). We found that prior optoFcR activation increased the amount of eating roughly 2-fold compared to cells that either did not receive prior light stimulation or did not express the optoFcR (Figure 2b). This demonstrates that prior FcR activation primes macrophages to respond to future IgG.

**Figure 2.**
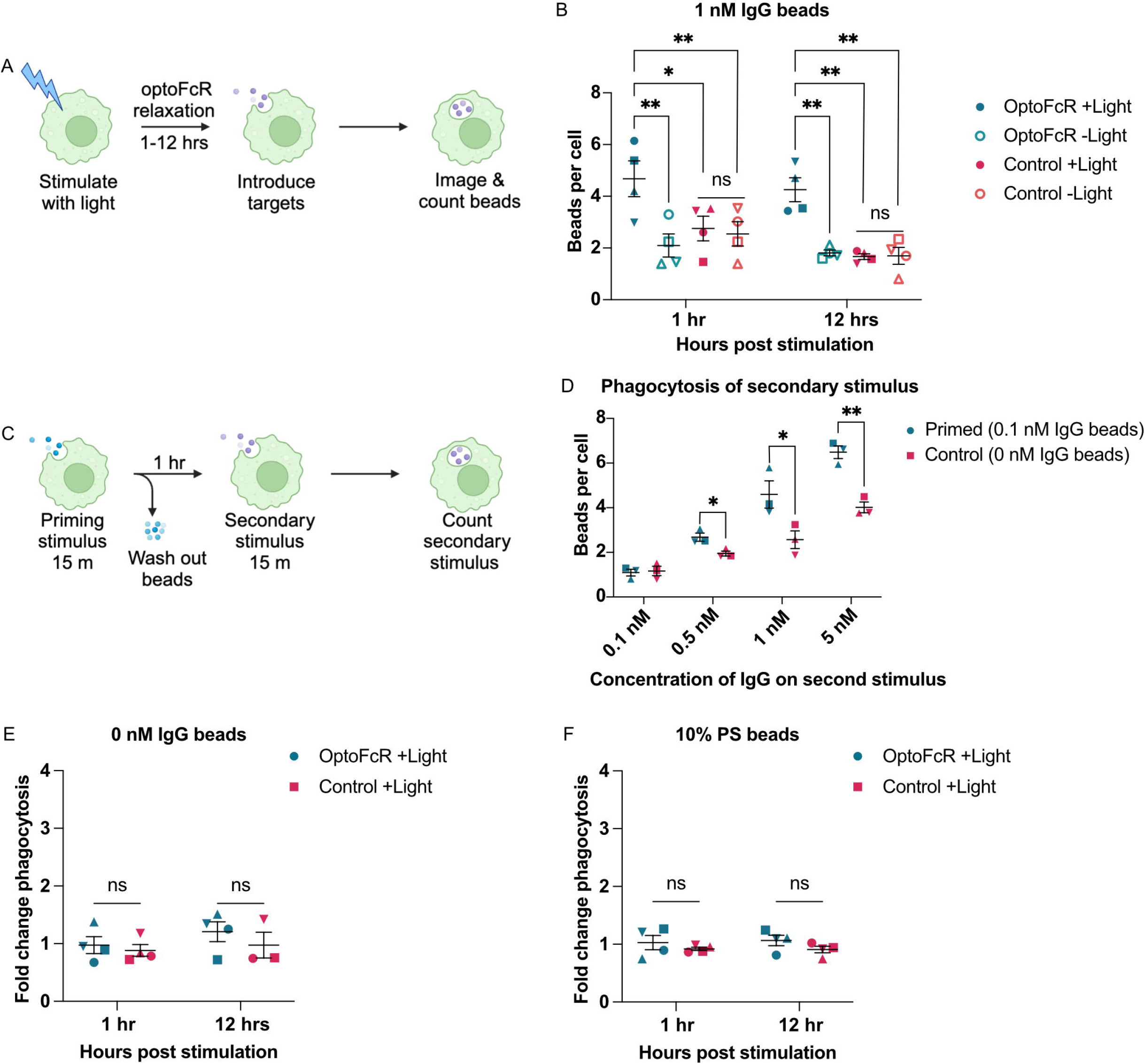
Prior FcR activation specifically enhances macrophage sensitivity to IgG. A) Schematic of experimental design for light-induced priming. Control (mCherry-CAAX) and optoFcR expressing BMDMs were stimulated with light for 15 min. The cells were returned to the dark for either 1 or 12 hrs, then targets opsonized with 1 nM IgG were introduced. After 15 minutes of phagocytosis, beads were washed out. Cells were imaged and the number of targets phagocytosed per cell was counted. B) Quantification of experiment described in (a). OptoFcR expressing cells that received light phagocytosed significantly more than cells that did not receive light, or cells that received light but did not express the optoFcR. C) Schematic of experimental design for priming macrophages with IgG beads. Wildtype BMDMs were given either beads with 0.1 nM or 0 nM IgG for 15 min as a priming stimulus. Those beads were then washed out and the cells were allowed to recover for 1 hr. A second set of different color beads ligated to IgG at the indicated concentration were added and the amount of phagocytosis was determined after 15 min. D) Quantification of phagocytosis for experiment described in (c). Macrophages that were preincubated with 0.1 nM IgG-conjugated beads phagocytosed significantly more than macrophages preincubated with unopsonized beads. E) Phagocytosis of bead targets without an ‘eat-me’ signal. Experiment was performed as described in (a) except the beads did not contain IgG. Phagocytosis is normalized to unstimulated control cells. F) Phagocytosis of bead targets with the efferocytic ‘eat-me’ signal, phosphatidylserine. Phagocytosis is normalized to unstimulated control cells. Experiment was performed as described in (a) except the beads were covered in a 10% phosphatidylserine bilayer to mimic an apoptotic corpse. In all graphs, each point represents the mean of an independent experiment. Data collected in the same replicate are denoted by symbol shape. Bars represent the mean and SEM. * indicates p<0.05, **indicates p<0.005, **** indicates p<0.0001 by two way anova with Sidak corrections (b), multiple T-tests with Holm-Sidak corrections (d), and unpaired t-test (e,f). See also supplemental figure 2.

Next, we systematically varied the light intensity and stimulation time used to activate the optoFcR as well as the delay between activating the optoFcR and measuring phagocytosis of IgG coated beads. In all cases, we found that low doses of light, and thus less FcR activity, led to the highest macrophage priming (Figure S2). The amount of light that best primed macrophages was not sufficient to activate phagocytosis on its own (Figure 1h). This suggests that a sub-threshold level of FcR activation, not sufficient to activate phagocytosis, primes macrophages for future encounters with IgG.

We then wanted to know if we could induce macrophage priming using IgG and the endogenous FcR. Based on our data from the optoFcR, we predicted that a low level of IgG exposure would enhance phagocytosis of a second dose of IgG-coated beads. To test this, we preincubated macrophages with beads containing an IgG density that did not activate phagocytosis (Figure S1) or unopsonized beads. We then extensively washed the macrophages to remove these beads and added a second color of IgG-bound beads (Figure 2c). We found that macrophages preincubated with IgG-coated beads phagocytosed more than macrophages pre-incubated with unopsonized beads when the second dose was above the threshold for inducing phagocytosis (Figure 2d). This suggests that the native IgG and FcR system primes macrophages like the optoFcR system.

### Prior FcR activation specifically enhances phagocytosis of IgG coated beads, not phosphatidylserine or unopsonized beads

We next wanted to know if prior FcR activation specifically primed macrophages to phagocytose IgG-coated targets or if it broadly enhanced phagocytosis. We first determined if prior optoFcR activation increased non-specific phagocytosis of unopsonized targets. Primed macrophages did not phagocytose more unopsonized beads than control macrophages (Figure 2e). We then determined if prior optoFcR activation increased efferocytosis, the engulfment of apoptotic cells. The molecular regulators of efferocytosis partially overlap with antibody-dependent phagocytosis, but the processes require some unique signaling pathways including different ‘eat-me’ signal receptors^26^. By integrating phosphatidylserine (PS), an efferocytic ‘eat-me’ signal, into the lipid mixture on our silica bead targets, we can recapitulate apoptotic corpse engulfment *in vitro*^27^. We incubated PS-coated beads with macrophages at 1 and 12 hours post light stimulation and measured the amount of eating. There was no change in the amount of eating at either timepoint (Figure 2f). These data suggest that prior FcR activation primes macrophages to specifically react to IgG. This also suggests that the molecular regulators of priming are unique to FcR signaling, rather than in one of the pathways shared by efferocytosis and antibody-dependent phagocytosis.

We then investigated if prior FcR stimulation changed macrophage sensitivity for IgG, lowering the threshold of IgG required for initiating phagocytosis. Alternatively, priming could increase macrophage capacity, the maximum number of targets each macrophage can engulf. To do this, we added beads with various concentrations of IgG to BMDMs and calculated the phagocytic index (Figure S2). Primed macrophages show enhanced eating of beads with low concentration of IgG. The total capacity for phagocytosis is not significantly changed in primed macrophages. This indicates that priming primarily alters macrophage sensitivity to low levels of IgG, rather than overall capacity for phagocytosis.

### Primed macrophages phagocytose more antibody opsonized cancer cells

We next sought to determine if primed macrophages can increase whole cell eating of opsonized cancer cell targets. In addition to phagocytosis, macrophages often trogocytose or nibble target cells, stripping the cancer cells of target antigen without killing them^15^. Prior clinical studies have shown that the anti-CD20 therapeutic antibody (rituximab) is most effective when administered more frequently at a low dose, which minimizes antigen shaving^28^. Given our results, we hypothesized that prior FcR activation might also enhance phagocytosis of cancer cells. We first measured phagocytosis of Raji B cells incubated with increasing concentrations of anti-CD20 antibody to find an antibody concentration where we could detect a change in macrophage sensitivity (Figure S3). To measure both the amount of whole cell phagocytosis and trogocytosis, we incubated IgG opsonized Raji cell targets with primed or unprimed optoFcR-expressing BMDMs and analyzed the cells with timelapse microscopy (Figure 3a-b; Video S4-6). We found that the number of Raji cells phagocytosed, and the percent of phagocytic macrophages increased in primed macrophages (Figure 3c-d). The percentage of primed macrophages that trogocytosed was not significantly increased compared to unprimed macrophages (Figure 3e). This suggests that primed macrophages are better at phagocytosing antibody-opsonized cancer cells.

**Figure 3.**
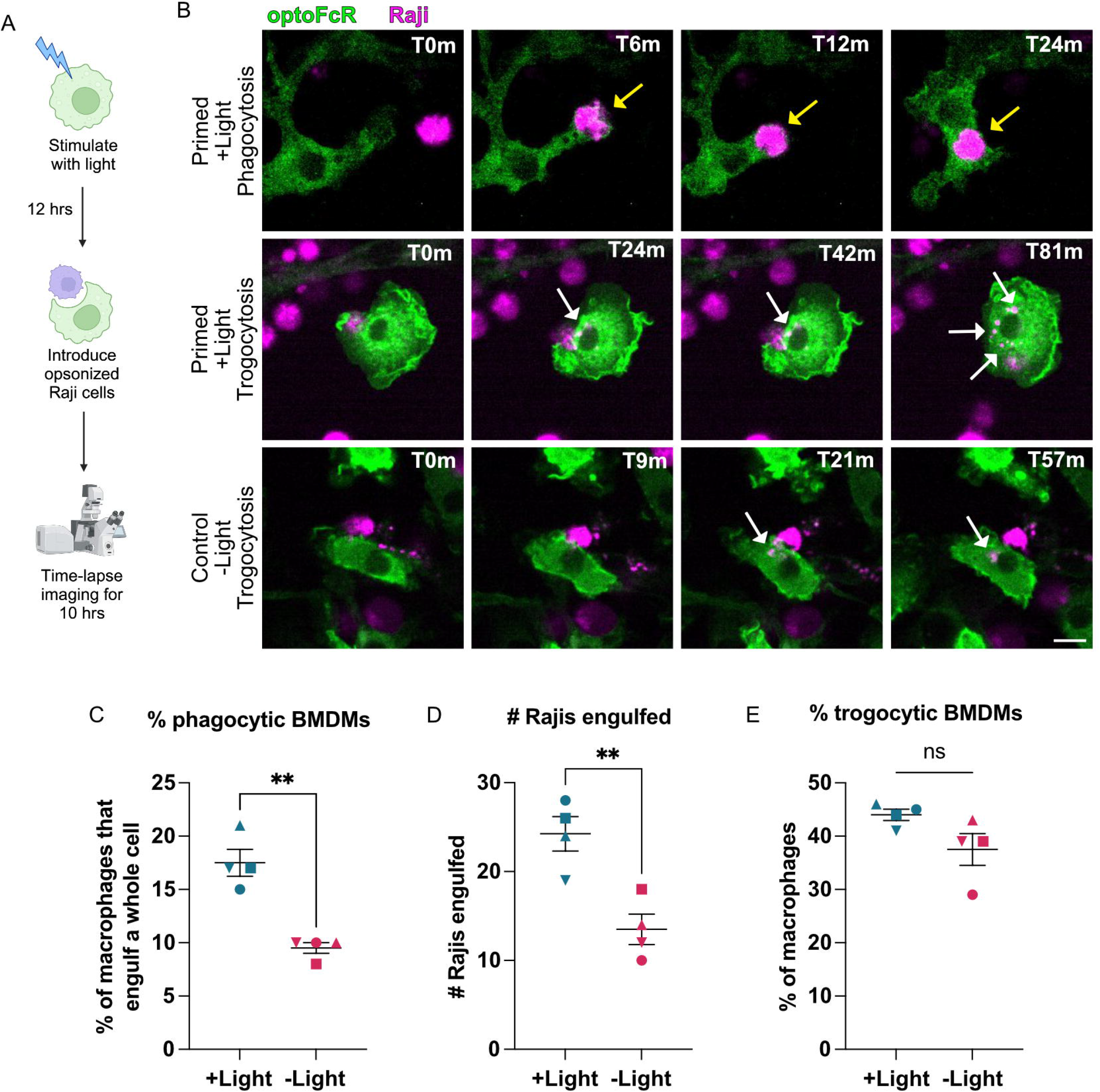
Macrophages are primed for increased cancer cell engulfment. A) Schematic of experimental design for cancer cell targets. optoFcR cells were exposed to light for 15 minutes, then returned to the dark. Raji B cells opsonized with 5 ng/ml IgG were added after 12 hours, and the cancer cell-macrophage interactions were captured by timelapse microscopy. B) Representative images of optoFcR expressing macrophages completing phagocytosis and trogocytosis. Raji B cells were visualized using cell trace far red (magenta) and macrophages were visualized using optoFcR-mSc (green). Yellow arrows point to phagocytosis of a whole cancer cell and white arrows point to trogocytosis of cancer cell fragments. See also supplemental movies 4-C) Percent of BMDMs that engulfed a whole target cell increased with prior FcR stimulation. D) The number of whole raji cells engulfed was greater in primed macrophages. E) Percent of BMDMs that trogocytosed a raji target was not significantly different with priming. Each data point represents the mean of an independent experiment, denoted by symbol shape, and bars represent the mean and SEM. **indicates p<0.005 using an unpaired t-test. See also supplemental figure 3.

### Macrophage priming occurs through a short-term and long-term mechanism

We next sought to determine the molecular mechanism for enhanced phagocytosis after FcR activation. We observed enhanced phagocytosis at both 1 hour and 12 hours after optoFcR stimulation (Figure 2b). While 12 hours post-stimulation is likely enough time for changes in transcription or translation to affect macrophage phenotypes, 1 hour is likely too short for this mechanism. We decided to carefully assay when macrophage priming occurred. To do this we activated the optoFcR and varied the time before presenting a second stimulus of IgG coated beads and measuring phagocytosis. We saw robust priming occurring in two discrete waves: a short-term response that peaks around 1 hour after FcR activation, and a long-term response that begins at 4 hours after FcR activation and persists for at least 24 hours (Figure 4a).

**Figure 4.**
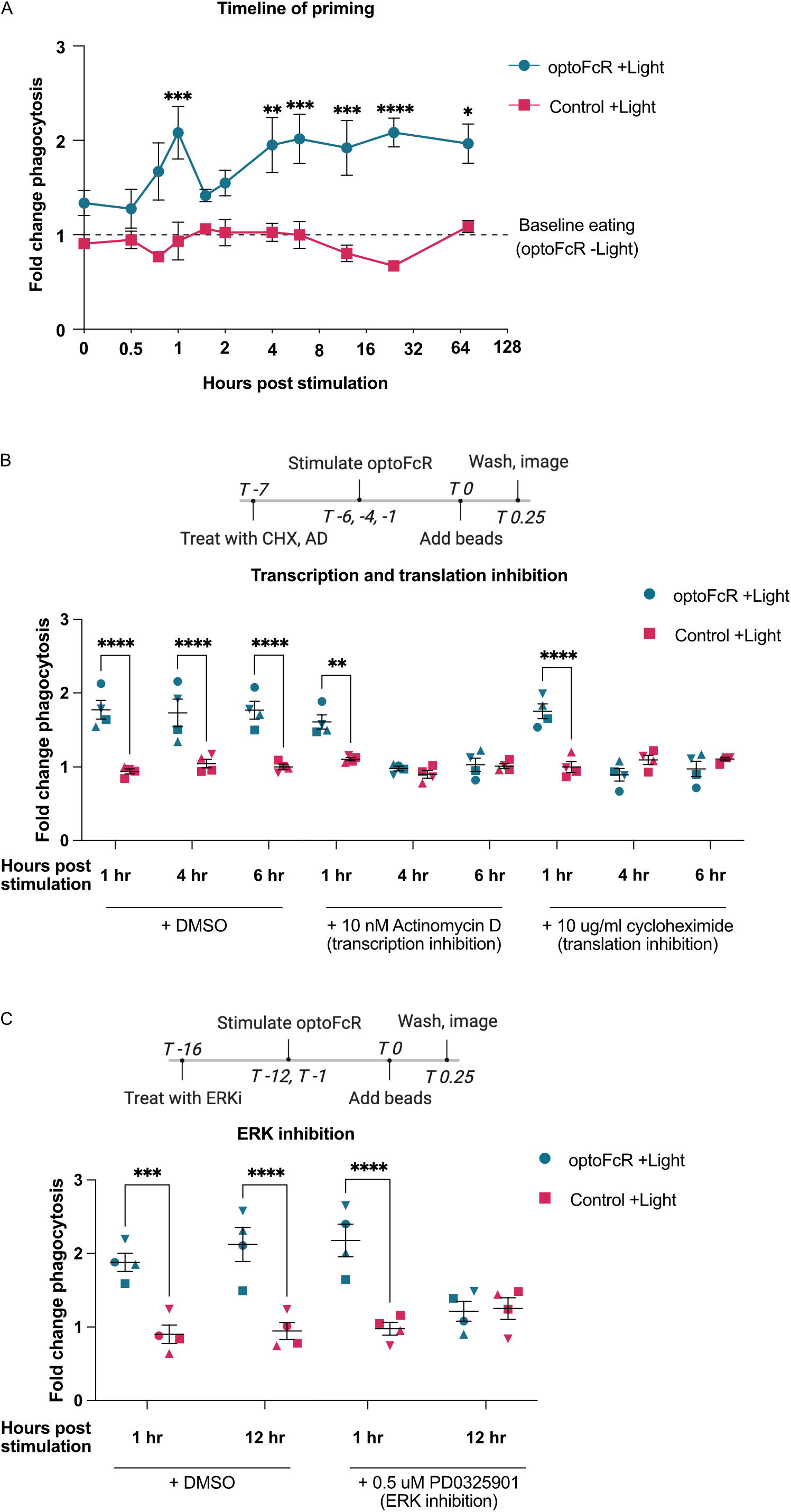
FcR mediated priming occurs via a short- and long-term mechanism. A) Phagocytosis of 1nM IgG beads added at the indicated time post 15 min low intensity light stimulation. Blue line denotes macrophages expressing the optoFcR and red denotes macrophages expressing a control mCh-CAAX. Enhanced phagocytosis occurs in two discrete peaks and lasts for at least 72 hrs. Points denote the mean of 4 independent replicates. Phagocytosis is normalized to unstimulated control cells. B) Macrophages were treated with Actonomycin D (AD, 10nM) or Cycloheximide (CHX, 10ug/ml) to block transcription or translation respectively starting 7 hours before phagocytosis. Macrophages were stimulated with light to activate the optoFcR at 1, 4 or 6 hours before phagocytosis. AD and CHX did not eliminate priming at 1 hour post-stimulation. AD and CHX eliminated the enhanced phagocytosis phenotype at 4 and 6 hours post stimulation. C) Macrophages were treated with Erk inhibitor (PD0325901, 0.5uM) or DMSO control for 16 hrs before measuring phagocytosis. optoFcR macrophages stimulated with light 1 hour before bead addition still showed enhanced phagocytosis. Macrophages stimulated with light 12 hours before bead addition phagocytosed the same number of beads as controls. Phagocytosis is normalized to unstimulated control cells. Each data point represents the mean of an independent experiment. Data collected in the same replicate are denoted by symbol shape. Bars represent the mean and SEM. *indicates p<0.05, **indicates p<0.005, ***indicates p<0.0005, **** indicates p<0.0001 using a two way anova with Sidak corrections (a-c).

As the long-term priming following optoFcR activation persists for at least 72 hours, we speculated that this response requires *de novo* protein production rather than a more transient post-translational modification mechanism. We evaluated priming in macrophages treated with cycloheximide (CHX) and actinomycin D (AD) to inhibit translation and transcription respectively. Treatment with either CHX or AD significantly reduced phagocytosis in primed macrophages compared to DMSO treated control macrophages at 4 and 6 hours post light stimulation (Figure 4b). This suggests that *de novo* mRNA and protein synthesis is required for a long-term memory response. Blocking new protein synthesis did not significantly reduce priming at 1 hour post stimulation, suggesting that short-term memory is not reliant on new protein production (Figure 4b). Overall this suggests that there are two mechanisms for macrophage priming – one that operates on a short timescale and does not require synthesis of new proteins, and one that operates on a long time scale and requires changes in gene expression.

### Erk activation is required for long-term priming

Because long term priming requires new protein production, we sought to dissect which transcriptional programs were being executed by the macrophages. Erk, a nuclear kinase, functions downstream of the FcR and regulates the macrophage dose dependent response to LPS as well as many other immune signaling pathways^11^. To determine if Erk contributes to macrophage priming, we used PD0325901 to block Erk activity. We then stimulated the optoFcR for 15 minutes, waited 1 or 12 hours, and measured phagocytosis of IgG-coated beads. Inhibiting Erk signaling blocked long-term memory with no effect on short-term memory (Figure 4c). These results indicate that long-term priming requires a transcriptional response mediated by Erk activation.

### Initiation of engulfment proceeds faster and has a higher success rate in primed macrophages

Having demonstrated that short-term priming does not require new protein synthesis like long-term priming, we sought to determine the mechanism for short term priming. To do this, we first wanted to isolate which step in the phagocytic process was enhanced by prior sub-threshold activation of the FcR. We quantified the kinetics of engulfment using live cell imaging, breaking the process of phagocytosis into three steps: target binding, initiation of phagocytosis, and completion^21^ (Figure 5a,b; Video S7). We then quantified the time between each step in primed and unprimed macrophages, and the percent of bead contacts that successfully progressed from one step to the next without membrane retraction and target release. Primed macrophages were more likely to initiate phagocytosis of bound beads, and the time between bead binding and initiation was less (Figure 5c,d). In contrast, after initiation the chance of successfully completing phagocytosis and the speed of phagocytosis were the same in primed and unprimed macrophages (Figure 5e,f). Overall, the percent of bead contacts that result in successful phagocytic events is significantly increased in primed macrophages (Figure 5g). Stimulated control macrophages that do not express the optoFcR did not show a difference in any measure of phagocytosis compared to unstimulated macrophages (Figure S4). These data indicate that prior sub-threshold FcR activation primes macrophages for faster target recognition and more frequent signal initiation, implicating early phagocytic machinery.

**Figure 5.**
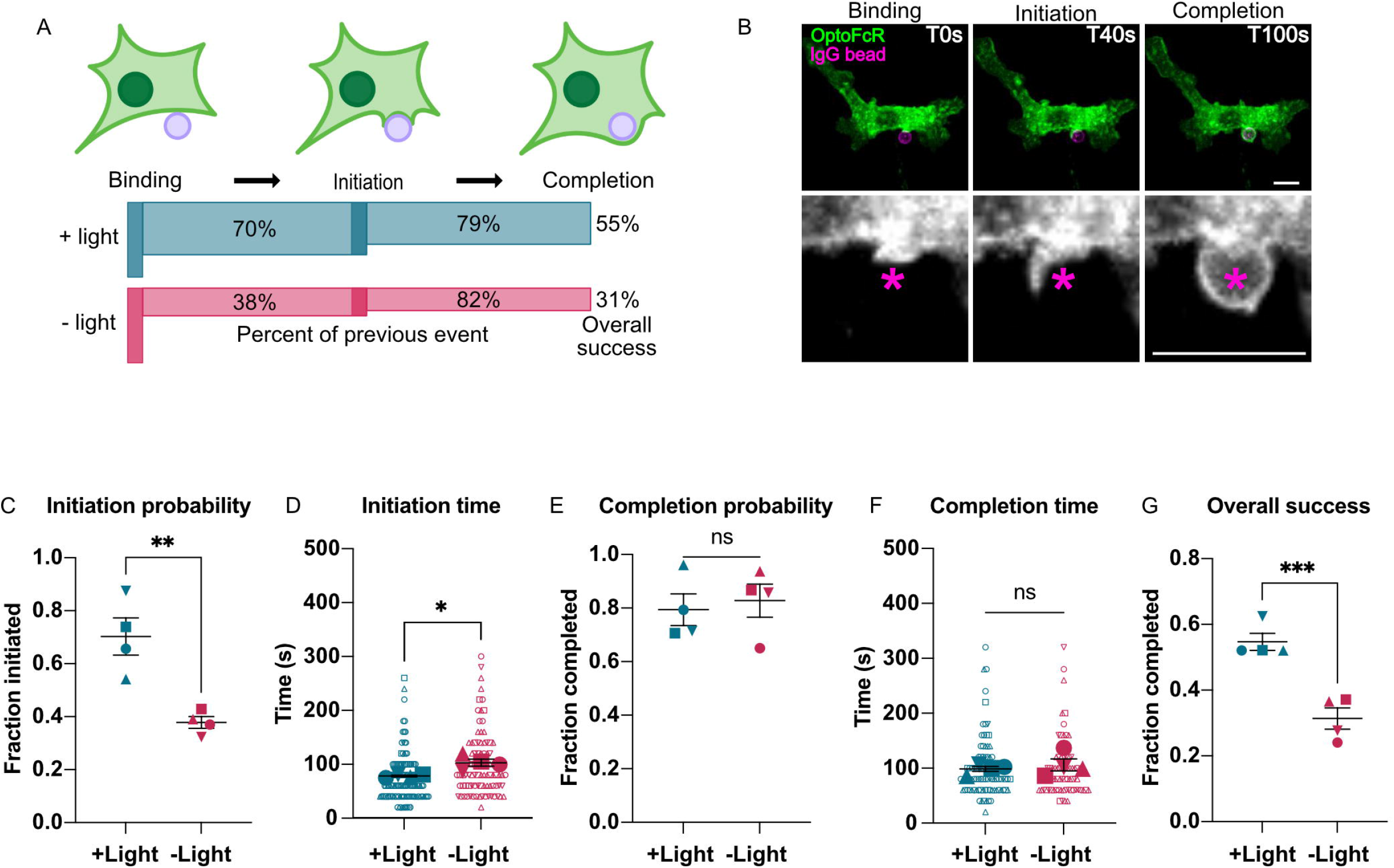
Initiation of phagocytosis is faster and more likely in primed macrophages. A) optoFcR macrophages received a 15 min light stimulus 1 hour before bead addition (primed) and were compared to optoFcR macrophages that did not receive light stimulation (unprimed). Timelapse imaging was used to quantify the kinetics of phagocytosis. Schematic shows each step in phagocytosis: binding (target contact with cell), initiation (formation of the phagocytic cup), and completion (cup closure and bead internalization). Below, the schematic shows percent of macrophage-bead contacts that progress from one stage to the next for primed and unprimed macrophages. B) Representative images of each stage. Scale bar is 10 um. See also supplemental movie 7 C) Percent of bead contacts that initiate phagocytosis is higher in primed macrophages. D) Time from binding to initiation in optoFcR primed macrophages is decreased compared with unprimed macrophages. E) Probability of completing phagocytosis after initiation is comparable in primed and unprimed macrophages. F) The speed of cup closure is comparable in primed and unprimed macrophages. G) The overall success rate of phagocytosis is higher in primed macrophages compared to unprimed macrophages. Each filled data point represents the mean of an independent experiment, denoted by symbol shape, and corresponding outlined data points represent individual bead times. Bars represent the mean and SEM. *indicates p<0.05, **indicates p<0.005, ***indicates p<0.0005 using an unpaired t-test on the means of each replicate. See also supplemental figure 4.

### Priming is associated with enhanced FcR mobility

As initiation of phagocytosis is faster in primed macrophages, and our prior data indicated that the molecular regulator of priming was specific to the FcR pathway, we decided to look at FcR mobility. Clustering of IgG, and subsequently FcR, increases the frequency and speed of initiating phagocytosis but not the speed of cup closure, similar to the phenotype we observed in our phagocytosis kinetics analysis^21^. FcR cluster formation and subsequent activation is dependent on the lateral mobility of FcRs, which is constrained by a heterogeneous F-actin ‘fence’ and other mechanisms^29–32^. Increased FcR mobility correlates with increased binding of IgG-coated targets and phagocytosis^29,31^. We hypothesized that primed macrophages may have higher FcR mobility, which could explain the increased speed and frequency of initiating phagocytosis. To determine if optoFcR priming increases receptor mobility, we tracked single FcR molecules on optoFcR primed and unprimed cells (Figure 6a). On average, the FcR molecules on primed macrophages had a higher mean jump distance (MJD; average distance traveled between frames) than on unprimed macrophages (Figure 6b). Graphing the MJD of individual tracks, primed macrophages had a multi-modal distribution of track MJDs, suggesting there may be a more mobile population in the primed macrophages (Figure 6c).

**Figure 6.**
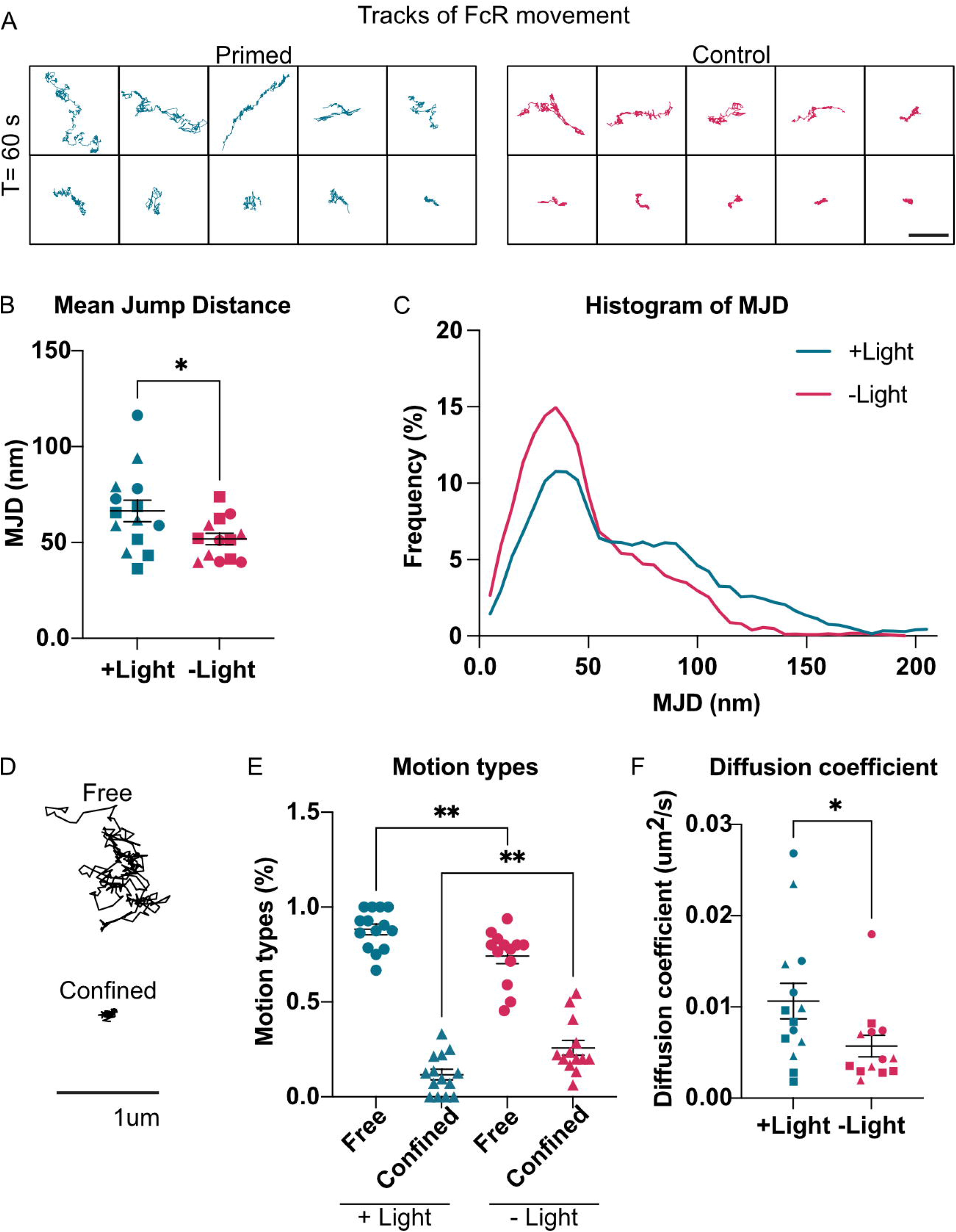
Priming increases FcR mobility. A) optoFcR macrophages were primed with a 15 minute light stimulus or were not stimulated. One hour later, single FcR molecules were labeled with QDots and tracked for 60 seconds. Tracks from a representative primed and unprimed image are shown. Scale bar is 1 um. B) Quantification of the per image mean jump distance (MJD) of all tracks. FcRs from primed macrophages had increased MJD compared to unprimed macrophages. C) Histogram of MJD from all acquired tracks from primed and unprimed macrophages. Primed macrophages show a population of FcRs that have a greater MJD than unprimed macrophages. Histogram was smoothed in prism using a second order polynomial with 6 neighbors on each size. D) Examples of ‘free’ and ‘confined’ tracks. E) Quantification of motion type analysis shows the proportion of tracks categorized as ‘free’ (circles) or ‘confined’ (triangles) for individual images. The percent of tracks defined as ‘free’ increases in optoFcR expressing macrophages exposed to light is compared to optoFcR macrophages with no light exposure. F) Quantification of the per image mean diffusion coefficients. Tracks from primed macrophages have a higher diffusion coefficient compared to unprimed macrophages. Each filled data point represents the mean from a single image. Images were acquired in three separate experiments, denoted by symbol shape (b and e). Bars represent the mean and SEM. *indicates p<0.05, **indicates p<0.005 using an unpaired t-test (b and f) or a one-way anova (e).

Previous studies have described FcR motion as free or confined^29,31^. Confined FcRs have limited mobility and may not be available to form signaling microclusters required for phagocytosis. To see if primed macrophages had more free FcRs, we categorized FcR tracks by motion type using a moment scaling spectrum analysis. Consistent with previous studies, we found a population of confined Fc Receptors that were restricted to small microdomains or ‘corals’ within the membrane (Figure 6d,e). This population was decreased in primed macrophages, suggesting that more FcRs may be available to join signaling microclusters. In addition, the average diffusion coefficient, indicative of how far each receptor travels, was significantly higher in primed macrophages (Figure 6f). Overall, this data suggests that FcRs are more mobile on primed macrophages, which may increase formation of FcR signaling clusters and the probability of initiating phagocytosis.

## Discussion

Our work suggests that the macrophage response to IgG integrates information from sequential encounters with IgG coated targets. Prior FcR activation makes macrophages more likely to phagocytose an IgG-coated target. This enhanced phagocytosis correlates with an increase in FcR mobility in the short term. Long term priming requires Erk signaling and transcription of new proteins.

Clustering of the FcR has previously been linked to its activation and enhanced phagocytosis^21,22,33,34^. We recapitulate this clustering using an optogenetic method and find that clustering of the optogenetic FcR is sufficient to drive Syk recruitment, which is indicative of phosphorylation of the intracellular Immunoreceptor Tyrosine-based Activation Motifs (ITAMs). The FcR has no inherent kinase activity, so why does clustering promote receptor activation? ITAM phosphorylation is controlled by the opposing actions of Src family kinases, which favor activation, and transmembrane phosphatases like CD45 that deactivate the receptor^1^. Prior work has shown that particle binding drives CD45 exclusion from the phagocytic synapse because the bulky extracellular domain is sterically excluded from the tight membrane-membrane interface^35–38^. This allows Src family kinases to dominate, tipping the kinase-phosphatase balance in favor of FcR activation^39^. Our prior work has shown that FcR clustering enhances ITAM phosphorylation independent of avidity effects that would increase receptor binding to the target^21^. This may be mediated by the formation of lipid ordered domains that segregate kinases and phosphatases, favoring immune receptor activation^40,41^. The optoFcR is tethered to the membrane via a myristoylation motif, favoring inclusion in lipid ordered domains, which may promote activation by Src family kinases and segregation from CD45 without the need for particle binding. Alternatively, the geometry of the clustered intracellular FcR domains may promote activation through another mechanism.

In the hour following optoFcR activation, the mobility of native FcRs on the macrophage surface increases. FcR mobility is constrained by the actin cytoskeleton and actin-associated transmembrane pickets^29,31,32,42^. Syk kinase activation rearranges the actin cytoskeleton allowing for less constrained FcR diffusion and greater mobility^31^. Since the optoFcR also recruits Syk, this may underlie the change in FcR mobility in our system. Our working model is that increased FcR mobility increases the formation of receptor clusters that promote initiating and continuing phagocytosis. This could enhance macrophage sensitivity by enabling formation of signaling clusters with a lower overall IgG density.

Prior work has shown a critical threshold of FcR activation is required for macrophages to commit to phagocytosis^10^. FcR signaling that is below this threshold activates the initial steps in the phagocytic signaling pathway, including recruitment of the downstream effector kinase Syk^10^. Whether this low level of activation has any effect on macrophages was not clear. Our data suggests that this low level of activation may have a purpose in the macrophage, allowing the cell to prepare for future encounters with IgG-bound targets.

Our data show that lower levels of optoFcR stimulation prime phagocytosis better than high levels. Prior work has shown that low levels of TLR activation prime macrophages for a rapid and strong response to future stimuli, without activating an inflammatory response alone^11^. Our study suggests this may be true for the FcR pathway as well. The effect of high levels of FcR activation may be different because it is associated with receptor internalization, which can lead to decreased phagocytic capacity^18^. Previous studies have shown that phagocytosing many antibody-coated cancer cells causes macrophage ‘hypophagia’ or reduced phagocytosis. This suggests that the signaling consequences of successful phagocytosis and sub-threshold FcR activation may be quite different.

While the adaptive immune system is traditionally thought of as the source of immunological memory, a growing body of evidence shows that the innate immune system also remembers prior infections and threats. This is often called “trained immunity” and occurs via epigenetic reprogramming of myeloid cells to increase or decrease their transcriptional response to reinfection^16,43,44^. Trained immunity can persist for years if myeloid progenitor cells are affected. Most of the work on trained immunity has focused on a memory of pathogenic molecules or inflammatory cytokines. Our work builds on this, suggesting that FcR activation also elicits a long term molecular memory. In contrast, the short term priming we describe is distinct from prior descriptions of trained immunity since it does not involve changes in gene expression.

The FcR is required for the full efficacy of many cancer immunotherapies, including popular immunotherapies like PD-1 and CTLA-4 blockades^9,45^. Some therapies, like the anti-CD20 antibody Rituximab, heavily rely on antibody-dependent cellular phagocytosis as a mechanism for eliminating cancer cells^4^. Interestingly, more frequent low dose treatments of Rituximab are more effective at treating Chronic Lymphocytic Leukemia (CLL) patients than higher dose treatments^28^. A key reason for this dosing schedule is to mitigate antigen shaving, or trogocytosis of target antigen. Enhancing phagocytosis without increasing antigen shaving is important but difficult. Our data shows that primed macrophages are better at phagocytosing whole cancer cells, but equally likely to trogocytose. This suggests that the current dosing regimen of frequent, low doses may already benefit from the effects of macrophage priming. Additionally, monocytes expressing Chimeric Antigen Receptors that signal through the FcR intracellular signaling domain are an exciting new avenue of cancer research^46–49^. How can we engineer hungrier macrophages to attack cancer cells? Our studies reveal that macrophage priming could enhance phagocytosis or other anti-cancer signaling pathways in these macrophages.

## Supporting information

Key Resources

## Acknowledgements

We thank members of the Morrissey lab for critical feedback on this manuscript. This work was supported by the UCSB Academic Senate, the National Institute of General Medical Sciences of the National Institutes of Health (R35 GM146935) and the UC Cancer Research Coordinating Committee (C23CR5592) to M.A.M. and the Eunice Kennedy Shriver National Institute of Child Health and Human Development of the National Institutes of Health (R01 HD108803-01) to M.Z.W. A.B. was supported by the Karl Storz Imaging fellowship. J.E.Q.N. was supported by a supplement to C23CR5592, the EUREKA scholars program and the MARC scholars program. A.G. was supported by the MARC scholars program. Schematics were created with BioRender.com. We thank the NRI-MCDB Microscopy Facility at UCSB, especially the director Ben Lopez for providing advice.

## Author contributions

Conceptualization, A.B., M.Z.W. and M.A.M.; Methodology, A.B., M.Z.W. and M.A.M.; Software, K.R. and S.J.R; Validation, A.B.; Formal Analysis, A.B., S.F., K.R. and S.J.R.; Investigation, A.B, S.F., E.N. and A.G.; Resources, A.B. and M.A.M.; Writing - Original Draft, A.B. and M.A.M.; Writing - Review and Editing, A.B. and M.A.M.; Visualization, A.B.; Supervision, M.A.M.; Funding Acquisition, M.A.M.

## Methods

### Resource Availability

#### Lead Contact

Further information and requests for resources and reagents should be directed to and will be fulfilled by the Lead Contact, Meghan Morrissey.

#### Bone-marrow derived macrophage cell culture

Six-to ten-week-old C57BL/6 mice were sacrificed by CO2 inhalation. Hips and femurs were dissected and bone marrow was harvested as described in Weischenfeldt and Porse^50^. Macrophage progenitors were differentiated for seven days in RPMI-1640, 10% FBS, 1% PSG supplemented with 20% L929-conditioned media. Macrophage differentiation was confirmed by flow cytometry identifying CD11b and F4/80 double positive cells. Differentiated BMDMs were used for experiments from days 7 to 11.

#### Lentivirus production and infection

All constructs were expressed in BMDMs using lentiviral infection. Lentivirus was produced in HEK293T cells transfected with pMD2.G (Gift from Didier Trono, Addgene plasmid # 12259 containing the VSV-G envelope protein), pCMV-dR8.2^51^ (Gift from Bob Weinberg, Addgene plasmid #8455), and a lentiviral backbone vector containing the construct of interest using lipofectamine LTX (Invitrogen, Catalog # 15338–100). The media was harvested 72 h post-transfection, filtered through a 0.45 μm filter (Millapore, Catalog #SLHVM33RS) and concentrated using LentiX (Takara Biosciences, Catalog #631232). Concentrated lentivirus was added to cells on day 2 of differentiation. Cells were analyzed between days 7-11.

#### Optogenetic stimulation

Cells receiving low intensity light (5 uW/cm^2^) and medium intensity light (190 uW/cm^2^) were stimulated using a LITOS LED illumination plate^52^ for 15 min. Cells receiving high intensity light were stimulated using the 488 laser at 75% laser power on a spinning disc confocal microscope for 1 s at 20 s intervals for a total of 15 m (1,389 uW/cm^2^). Intensity was determined using a slide power meter set to measure 450 nm light. Light used to prime cells was low intensity (5 uW/cm^2^) unless otherwise indicated.

#### Clustering and colocalization analysis

##### optoFcR clustering

50,000 BMDMs expressing the optoFcR were plated in one well of a 96-well glass bottom MatriPlate (Brooks, Catalog # MGB096-1-2-LG-L) between 12 and 24 h prior to the experiment. Cells were then continuously imaged with high intensity light stimulation for 30 min and then imaged for another 60 min without light stimulation. Clustering was determined by the presence of visible puncta in the cells.

##### SYK and optoFcR colocalization

50,000 RAW264.7 macrophages virally infected with both the optoFcR and SYK-NeonGreen (Addgene, Plasmid # 176610) were plated in one well of a 96-well glass bottom MatriPlate (Brooks, Catalog # MGB096-1-2-LG-L) between 12 and 24 h prior to the experiment. Cells were then continuously imaged with high intensity light stimulation for 30 min. Colocalization was determined by a pearsons correlation coefficient for the same membrane region at the first and last timepoints using the JaCoP plugin in ImageJ^53^.

#### ICAM-1 protein purification

ICAM-tagBFP-His_10_^54^ was expressed in SF9 or HiFive cells using the Bac-to-Bac baculovirus system as described previously^55^. Insect cell media containing secreted proteins was harvested 72 h after infection with baculovirus. His10 proteins were purified by using Ni-NTA agarose (QIAGEN, Catalog # 30230), followed by size exclusion chromatography using a Superdex 200 10/300 GL column (GE Healthcare, Catalog # 17517501). The purification buffer was 150 mM NaCl, 50 mM HEPES pH 7.4, 5% glycerol, 2 mM TCEP.

#### Supported lipid bilayer coated beads

##### SUV preparation

For IgG conjugated beads the following chloroform-suspended lipids were mixed and desiccated overnight to remove chloroform: 98.8% POPC (Avanti, Catalog # 850457), 1% biotinyl cap PE (Avanti, Catalog # 870273), 0.1% PEG5000-PE (Avanti, Catalog # 880230, and 0.1% atto390-DOPE (ATTO-TEC GmbH, Catalog # AD 390–161) or 0.1% atto647-DOPE (ATTO-TEC GmbH, Catalog # AD 647–161). The lipid sheets were resuspended in PBS, pH7.2 (GIBCO, Catalog # 20012050) at 10 mM concentration and stored under inert nitrogen gas.

For ICAM-1 conjugated beads, the following chloroform-suspended lipids were mixed and desiccated overnight to remove chloroform: 97.8% POPC (Avanti, Catalog # 850457), 2% DGS-NTA (Avanti, Catalog # 790404), 0.1% PEG5000-PE (Avanti, Catalog # 880230, and 0.1% atto390-DOPE (ATTO-TEC GmbH, Catalog # AD 390– 161). The lipid sheets were resuspended in PBS, pH7.2 (GIBCO, Catalog # 20012050) and stored under inert gas.

For PS beads the following chloroform-suspended lipids were mixed and desiccated overnight to remove chloroform: 89.8% POPC (Avanti, Catalog # 850457), 10% DOPS (Avanti, Catalog # 840035), 0.1% PEG5000-PE (Avanti, Catalog # 880230, and 0.1% atto390-DOPE (ATTO-TEC GmbH, Catalog # AD 390–161). The lipid sheets were resuspended in PBS, pH7.2 (GIBCO, Catalog # 20012050) and stored under inert gas.

For all SUVs, the lipids were broken into small unilamellar vesicles via several rounds of freeze-thaws. The lipids were then stored at-80°C under argon for up to six months. To remove aggregated lipids, the solution was diluted to 2 mM and filtered through a 0.22 uM filter (Millapore, Catalog # SLLG013SL) immediately prior to use.

##### Bead preparation

Silica beads with a 4.89 μm diameter (10% solids, Bangs Labs, Catalog # SS05003, Lot # 13427) were washed with PBS, mixed with 1mM SUVs in PBS and incubated at room temperature for 30 min with end-over-end mixing to allow for bilayer formation. Beads were then washed with PBS to remove excess SUVs and incubated in 0.2% casein (Sigma, Catalog # C5890) in PBS for 15 min before protein coupling (IgG and ICAM-1 beads). For IgG conjugated beads, anti-biotin AlexaFluor647-IgG (Jackson ImmunoResearch Laboratories Catalog # 200-602-211, Lot # 156182) was added at 1 nM to a 10x dilution of beads (1% solids), unless otherwise indicated. For ICAM-1 conjugated beads, ICAM-1 was added at 10nM. Proteins were coupled to the bilayer for 30 min at room temperature with end-over-end mixing.

##### Bead engulfment assay

50,000 BMDMs were plated in one well of a 96-well glass bottom MatriPlate (Brooks, Catalog # MGB096-1-2-LG-L) between 12 and 24 h prior to the experiment. ∼8 × 10^5^ beads were added to wells and engulfment was allowed to proceed for 15 min.

##### Inhibitors

For transcription and translation inhibited priming, 10 nM actinomycin D (Cell signaling, Catalog # 15021s) or 10 ug/ml cycloheximide (Cell signaling, Catalog # 2112s) were added to cells 7 hours prior to the start of the experiment. For ERK inhibited priming, 0.5 uM PD0325901 (Sigma, Catalog # PZ0162) was added to cells 16 hours prior to the start of the experiment.

##### Microscopy and analysis

Images were acquired on a spinning disc confocal microscope (Nikon Ti2-E inverted microscope with a Yokogawa CSU-W1 spinning disk unit and an Orca Fusion BT scMos camera) equipped with a 40 × 0.95 NA Plan Apo air and a 100 × 1.49 NA oil immersion objective. The microscope was controlled using Nikon Elements. Internalized particles were counted in ImageJ by a blinded analyzer using Blind-Analysis-Tools-1.0 ImageJ plug in.

#### Bead priming

50,000 BMDMs were plated in 1 well of a 96-well glass bottom plate 12-24 hrs prior to the start of the experiment. A priming dose of ∼8 × 10^5^ atto390 beads conjugated to either 1 nM or 0 nM IgG were added to the wells for 15 min. Any unengulfed beads were washed out 5x with media, then checked with a dissecting microscope to confirm the majority of engulfed beads had been removed. Then the cells were allowed to recover for 1 hour. ∼8 × 10^5^ atto647 beads prepared with the indicated IgG concentrations were then added to the wells and engulfment was allowed to proceed for 15 min. Cells were dyed with CellTrace CSFE (Thermo, Catalog # C34570) imaged and the number of atto647 beads engulfed per cell were counted.

#### Raji eating assay

40,000 BMDMs were plated in 1 well of a 96-well glass bottom plate 24 hrs prior to the experiment and stimulated with low intensity LITOS illumination 12 hrs prior to the experiment. Raji cells were dyed with CellTrace Far Red (Thermo, C34572), incubated with a human-mouse hybrid aCD20 (InvivoGen hcd20-mab10, 5 ng/ml), added to wells at 40,000 cells per well, and imaged immediately. 25 positions per well were automatically selected and imaged every 3 min for 10 hrs. Unless otherwise noted, 100 macrophages were randomly selected and scored by a blind analyzer. Phagocytic macrophages were characterized as BMDMs that engulfed 1 or more whole Raji cell targets. Trogocytic macrophages were characterized as BMDMs that engulfed portions of Raji targets. The number of Raji cells engulfed per 100 macrophages was also counted.

#### Kinetics of engulfment

BMDMs were plated as described in the bead engulfment assay 12-24 hrs prior to the experiment and stimulated with low intensity LITOS illumination 1 hr prior to the experiment. Using ND acquisition in Elements, 2-3 positions per well were manually selected. Approximately 4 x 10^5^ beads were added and phagocytosis was imaged at 20 s intervals through 7 z planes for 15 min. Only beads that bound within the first 12 min were counted.

#### Receptor labeling and single particle tracking

##### Fab generation

Fabs from rat anti-mouse FcR (Cell signaling, 101307) and rabbit anti-rat biotin conjugated (Invitrogen, 13-4813-85) were generated using the Pierce Fab Preparation Kit (Thermo, 44985) according to the manufacture’s protocol. In brief, antibodies were run through a Zeba desalting column before cleavage with immobilized papain for 3 hours with end over end mixing and digestion was confirmed via SDS-PAGE. Fab fragments were then purified using a NAb Protein A column.

##### Receptor labeling and imaging

Single FcRs were labeled as previously described^31^. In brief, cells were blocked for 5 min in RPMI supplemented with 5% goat serum. Then, cells were incubated with primary fab fragments for 10 min in blocking medium. Next, cells were incubated with biotinylated secondary fab fragments for 10 min. Finally, cells were washed with blocking media and incubated with streptavidin-congugated Qdot 655 (Thermo, Q10123MP) for 4 min and immediately imaged. Images were acquired at 10 fps for 1 min using ND acquisition in Elements on a spinning disc confocal microscope.

##### Tracking

Tracks and particle mean jump distance were generated using the trackmate plugin on ImageJ^56,57^. Tracks that were less than 50 frames or that contained gaps greater than 3 frames were discarded from analysis. Motion types and diffusion coefficients were determined using moment scaling spectrum analysis using the formula described in Ewers et al, 2005^31,58–60^.

#### Quantification and statistical analysis

Statistical analysis was performed in Prism 8 (GraphPad). The statistical test used is indicated in the relevant figure legend. Sample sizes were predetermined and indicated in the relevant figure legend.

