## Supplemental Figures for "Prior Fc Receptor activation primes macrophages for increased sensitivity to IgG via long term and short term mechanisms"

Supplemental Figure 1

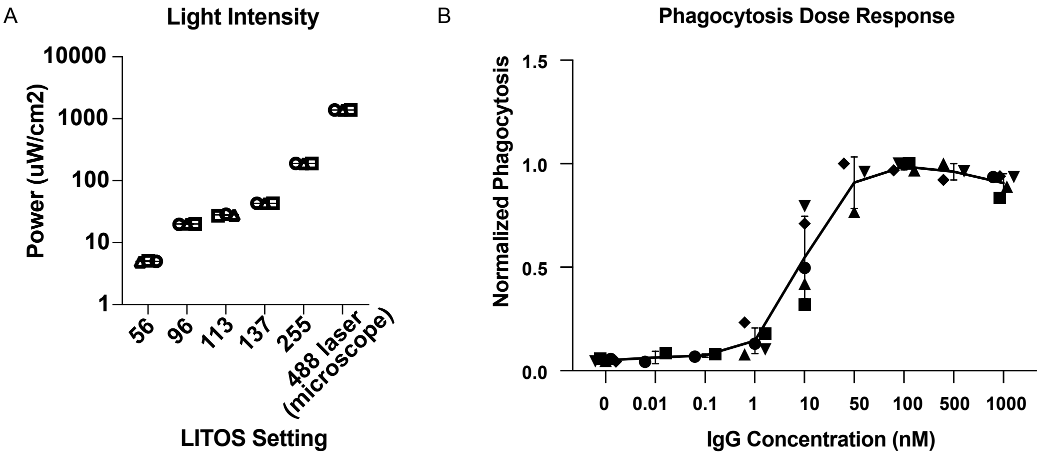


**Supplemental Figure 1; related to figure 1.**
A) Phagocytosis dose response curve for beads opsonized at the indicated concentrations of IgG in wildtype BMDMs. B) Slide power meter readouts from LITOS settings in a-b. Data are from 3 independent readings. All following experiments were performed with low intensity light (LITOS 56, 5 $uW/cm^2$) unless otherwise indicated. Medium (LITOS 255, 190 $uW/cm^2$) and high (488 confocal laser, 1390 $uW/cm^2$) used in Figure 1g are also depicted. Each data point represents the mean of an independent experiment, denoted by symbol shape, and bars represent the mean and SEM.

Supplemental Figure 2

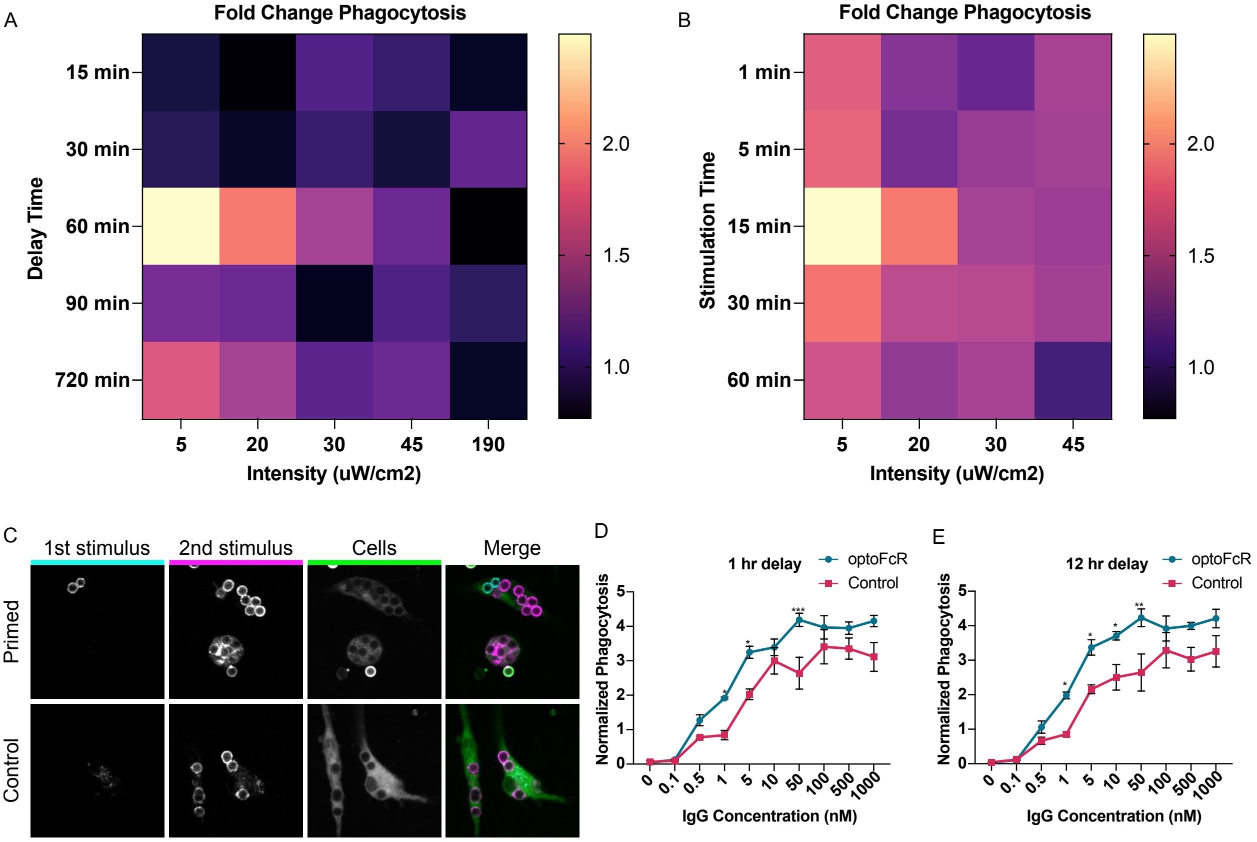


**Supplemental Figure 2; related to figure 2.**

A-B) Heat maps of fold change in phagocytosis of IgG opsonized beads in optoFcR expressing macrophages that received a priming dose of light compared to optoFcR expressing macrophages that did not receive light stimulation. In (a) the intensity and delay time were altered. In (b) the priming dose of light and stimulation time were altered. Data are from 3 independent replicates. C) Representative images of bead priming related to Figure 2d. First stimulus was 0.1nM IgG conjugated beads or unconjugated beads (atto390; cyan) for primed and control respectively. The second stimulus (atto647; magenta) was 1nM IgG conjugated beads for both conditions. D) Phagocytosis of beads opsonized with the indicated IgG concentration by optoFcR or control (CAAX-mCherry) expressing macrophages. Macrophages were stimulated with 15 min of low intensity light 1 hr before bead addition. Phagocytosis was normalized to phagocytosis of 1 nM IgG beads by unstimulated control cells. E) Phagocytosis of beads opsonized with the indicated IgG concentration by optoFcR or control (caax-mCherry) expressing macrophages. Macrophages were stimulated with 15 min of low intensity light 12 hrs before bead addition. Phagocytosis was normalized to phagocytosis of 1 nM IgG beads by unstimulated control cells. Each data point represents the mean of 4 independent experiments and bars represent SEM. *indicates $p<0.05$, **indicates $p<0.005$, **** indicates $p<0.0001$ using two way anova with Sidak corrections.

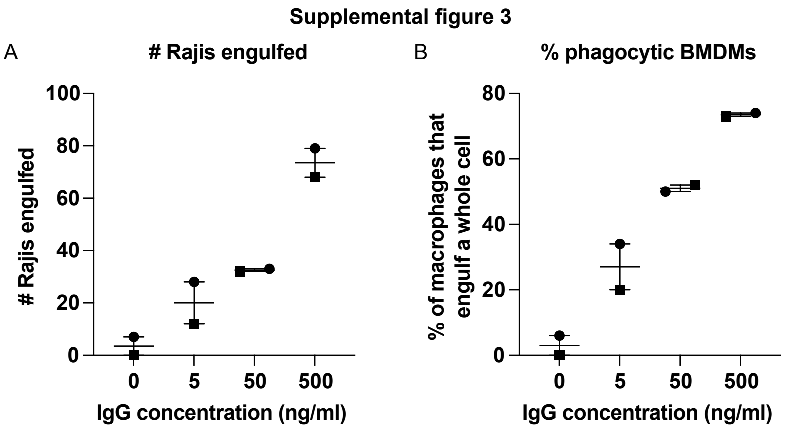


**Supplemental Figure 3; related to figure 3.**

A) Number of whole Raji cells engulfed by wildtype BMDMs at various concentrations of IgG as measured by timelapse microscopy. B) Percent of phagocytic wildtype BMDMs at various concentrations of IgG. Each data point represents the mean of an independent experiment in which 50 BMDMs were tracked for 10 hours, denoted by symbol shape, and bars represent the mean and range.

Supplemental figure 4

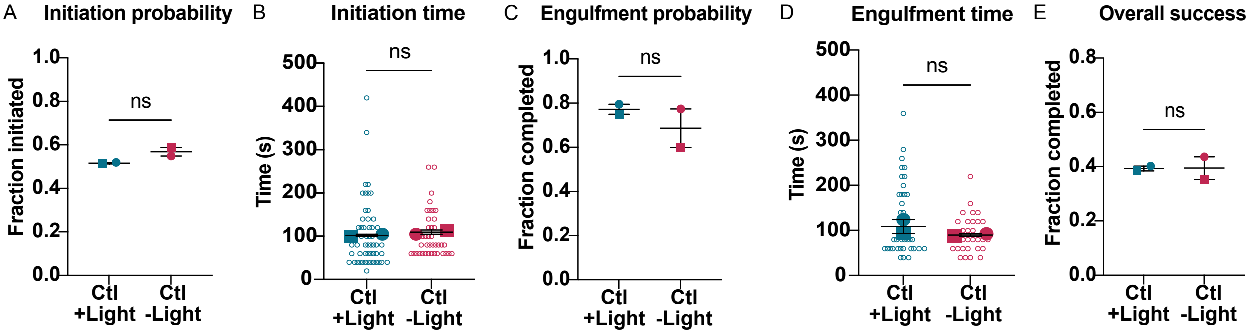


**Supplemental Figure 4; related to figure 5.**

A) Percent of bead contacts that initiate phagocytosis is unchanged in control cells (caax-mCherry) with and without light stimulation. D) Time from binding to initiation is unchanged in control cells with and without light stimulation. E) Completion rate of phagocytosis after initiation is unchanged in control cells with and without light stimulation. F) The speed of cup closure is unchanged in control cells with and without light stimulation. G) The overall success rate of phagocytosis is unchanged in control cells with and without light stimulation. Each filled data point represents the mean of an independent experiment, denoted by symbol shape, and corresponding outlined data points represent individual bead times. Bars represent the mean and range. Data was analyzed using an unpaired t-test.

**Video S1**
Representative movie of an optoFcR (visualized with mScarlet; white) expressing bone marrow derived macrophage stimulated with high intensity light. Light was on from 0-15 min and was turned off from 15-55 min. Images were taken every 30s.

**Video S2**
Representative movie of a RAW macrophages expressing the optoFcR (mScarlet; green) and SYK-neon (magenta) stimulated with high intensity light for the entire duration of the movie. Images were taken every 30s.

**Video S3**
Representative movie of an optoFcR (mScarlet; green) expressing bone marrow derived macrophage phagocytosing ICAM conjugated beads (atto390; magenta) with high intensity light stimulation. Light was on for the entire duration of the movie. Images were taken every 30s.

**Video S4**
Representative movie of a primed optoFcR (mScarlet; green) expressing macrophage phagocytosing a Raji cell target (CellTrace Far Red; magenta). The macrophage was stimulated with low intensity light for 15 minutes 12 hrs prior to the experiment. Images were taken every 3 min.

**Video S5**
Representative movie of a primed optoFcR (mScarlet; green) expressing macrophage (green) trogocytosing a Raji cell target (CellTrace Far Red; magenta). The macrophage was stimulated with low intensity light 12 hrs prior to the experiment. Images were taken every 3 min.

**Video S6**
Representative movie of an optoFcR (mScarlet; green) expressing macrophage trogocytosing a Raji cell target (CellTrace Far Red; magenta). The macrophage was not stimulated with light prior to the experiment. Images were taken every 3 min.

**Video S7**
Representative movie of an optoFcR expressing macrophage (caax-mCherry and optoFcR-mScarlet; green) phagocytosing a 1nM IgG conjugated bead (atto647; magenta). Movie was generated from a maximum projection of 7 z-stacks, 1.5um apart. Images were taken every 20s.
