## Supplementary material for "Prior Fc Receptor activation primes macrophages for increased sensitivity to IgG via long term and short term mechanisms": Key Resources

**Key resources table**

| **REAGENT or RESOURCE** | **SOURCE** | **IDENTIFIER** |
| --- | --- | --- |
| **Antibodies** | | |
| AlexaFlour647 anti-biotin IgG | Jackson ImmunoResearch Laboratories | Cat# 200-602-211, RRID:AB_2339046) |
| Anti-CD20 | InvivoGen | Cat# hcd20-mab10, RRID:AB_11124933 |
| Mouse anti-FcR | Cell signaling | Cat# 101307 |
| Anti-Rat IgG biotin conjugate | Invitrogen | Cat# 13-4813-85 |
| **Chemicals, peptides, and recombinant proteins** | | |
| POPC | Avanti | Cat# 850457 |
| Biotinyl cap PE | Avanti | Cat# 870273 |
| PEG5000-PE | Avanti | Cat# 880230 |
| Atto390-DOPE | ATTO-TEC GmbH | Cat# AD 390-161 |
| Atto647-DOPE | ATTO-TEC GmbH | Cat# AD 647-161 |
| Ni2+-DGS-NTA | Avanti | Cat# 790404 |
| DOPS | Avanti | Cat# 840035 |
| CellTrace Far Red | ThermoFisher | Cat# C34572 |
| CellTrace CSFE | ThermoFisher | Cat# C34570 |
| Qdot 655 | ThermoFisher | Cat# Q10123MP |
| Casein | Sigma | Cat# C5890 |
| PD0325901 - ERK inhibitor | Sigma | Cat# PZ0162 |
| Actinomycin D | Cell signaling | Cat# 15021s |
| Cycloheximide | Cell signaling | Cat# 2112s |
| **Critical commercial assays** | | |
| Pierce Fab Preparation Kit | ThermoFisher | Cat# 44985 |
| **Experimental models: Cell lines** | | |
| Human: HEK239T | ATCC | Cat# ATCC CRL-3216 |
| Mouse: RAW264.7 | ATCC | Cat# ATCC TIB-71; RRID:CVCL_0493 |
| Mouse: L929 | ATCC | Cat# ATCC CLL1; RRID:CVCL_0462 |
| Sf9 | ThermoFisher | Cat# 11496015 |
| **Experimental models: Organisms/strains** | | |
| Mouse: C57BL/6 | The Jackson Laboratory | Cat# 000664 |
| **Recombinant DNA** | | |
| pHR-optoFcR | This paper | In pHR vector. Myristolization sequence: MGSSKSKPKDPSQR; cytoplasmic domain (aa 45–86) of the Fc γ-chain UniProtKB - P20491 (FCERG_MOUSE); linker: STSG; fluorophore: mScarlet; linker: SDPGSGS; Cry2-olig (aa 1-498) of CRY2_ARATH UnitProtKB – Q96524 with E490G mutation followed by: ARDPP (as described in Taslimi et al, 2014). |
| pHR-Syk NeonGreen NK83 | Kern et al, 2021 | Addgene plasmid # 176610 ; http://n2t.net/addgene:176610 ; RRID:Addgene_176610 |
| pHR mCherry-caax | Morrissey et al, 2018 | mCherry fused to membrane targeting sequence from KRAB (amino acids LEKMSKDGKKKKKKSKTKCVIM) |
| pMD2.G | pMD2.G was a gift from Didier Trono | Addgene plasmid # 12259 ; http://n2t.net/addgene:12259 ; RRID:Addgene_12259 |
| pCMV-dR8.2 | pCMV-dR8.2 dvpr was a gift from Bob Weinberg | Addgene plasmid # 8455 ; http://n2t.net/addgene:8455 ; RRID:Addgene_8455 |
| ICAM-tagBFP-His10 | O’Donoghue et al 2013 | N/A |
| **Software and algorithms** | | |
| ImageJ -Fiji | NIH - https://fiji.sc/ | RRID:SCR_002285 |
| Affinity Designer | Serif | RRID:SCR_016952 |
| Prism | Graphpad | RRID:SCR_002798 |
| TrackMate | Ershov et al, 2022;  Tinevez et al, 2017 | N/A |
| Moment Scaling Spectrum analysis | This paper | N/A |
| Blind-Analysis-Tools-1.0 | Github https://github.com/ahtsaJ/Blind-Analysis-Tools | N/A |
| JaCoP | Bolte et al, 2006 | N/A |
| **Other** | | |
| 5 um silica beads | Bangs | Cat# SS05003 |
| MatriPlate | Brooks | Cat# MGB09-1-2-LG-L |
| LITOS stimulation plate | Hohener et al, 2022 | N/A |
